# Training Data Distribution Significantly Impacts the Estimation of Tissue Microstructure with Machine Learning

**DOI:** 10.1101/2021.04.13.439659

**Authors:** Noemi G. Gyori, Marco Palombo, Christopher A. Clark, Hui Zhang, Daniel C. Alexander

## Abstract

**Purpose:** Supervised machine learning (ML) provides a compelling alternative to traditional model fitting for parameter mapping in quantitative MRI. The aim of this work is to demonstrate and quantify the effect of different training strategies on the accuracy and precision of parameter estimates when supervised ML is used for fitting.

**Methods:** We fit a two-compartment biophysical model to diffusion measurements from in-vivo human brain, as well as simulated diffusion data, using both traditional model fitting and supervised ML. For supervised ML, we train several artificial neural networks, as well as random forest regressors, on different distributions of ground truth parameters. We compare the accuracy and precision of parameter estimates obtained from the different estimation approaches using synthetic test data.

**Results:** When the distribution of parameter combinations in the training set matches those observed in similar data sets, we observe high precision, but inaccurate estimates for atypical parameter combinations. In contrast, when training data is sampled uniformly from the entire plausible parameter space, estimates tend to be more accurate for atypical parameter combinations but may have lower precision for typical parameter combinations.

**Conclusion:** This work highlights the need to consider the choice of training data when deploying supervised ML for estimating microstructural metrics, as performance depends strongly on the training-set distribution. We show that high precision obtained using ML may mask strong bias, and visual assessment of the parameter maps is not sufficient for evaluating the quality of the estimates.

## 1. Introduction

Clinically used magnetic resonance imaging (MRI) typically focuses on the qualitative assessment of image contrast that arises from a combination of different properties of the imaged tissue, imaging hardware and measurement settings. Going a step further, quantitative MRI (qMRI) aims to quantify inherent tissue properties, such as T1- and T2-relaxation times, proton density, magnetisation transfer, susceptibility and diffusivity, by removing confounding effects arising from differences in imaging setup. Quantifying physical tissue features has many potential benefits, such as ease of interpretation, reproducibility, and straightforward comparisons between measurements made at different times or across different populations [1]. However, to quantify the tissue features of interest, it is necessary to define a model linking those features to the measured MRI signal and fit it to appropriately collected data. For example, in diffusion MRI (dMRI), a rich arsenal of biophysical models, signal representations and acquisition strategies have been proposed to quantify several tissue properties, such as mean diffusivity, microscopic anisotropy, neurite density and dispersion [2] [3]. One of the key challenges in qMRI is therefore estimating tissue features accurately, precisely and in a reproducible way, given a model and MRI data.

Conventionally, model fitting is performed voxel-by-voxel using optimisation techniques, often based on minimising a non-linear objective function. However, as models become more complex, conventional fitting approaches become slow and prone to local minima, and the estimation performance degrades with decreasing amount of available data and signal-to-noise ratio (SNR). These drawbacks can hamper the widespread use of qMRI in clinically relevant applications.

Recently, machine learning (ML) has emerged as a promising tool for overcoming many of the challenges associated with model fitting for qMRI. For example, ML methods based on artificial neural networks have been used to reduce estimation time of myelin water fraction in the brain [4] and to estimate T1 and T2 in a fast and robust way using sparse data from magnetic resonance fingerprinting [5]; whereas ML methods based on convolutional neural network approaches have been developed to estimate susceptibility using a single subject orientation [6]. In dMRI, ML has been used, for example, to bridge the gap between data-hungry imaging techniques and clinically feasible scans, for example by reconstructing super-resolved maps from low spatial resolution data [7] [8], or by estimating advanced diffusion-based metrics from sparse q-space acquisitions [9] [10] [11].

Most of the ML methods used in qMRI are based on the so-called supervised learning paradigm which relies on learning patterns from large amounts of examples, or training data, to map inputs to desired outputs. A key issue with supervised ML is that in the absence of balanced training data, the ML model may learn disruptive patterns. There are compelling examples of this in healthcare technology, where racial [12] and gender [13] biases arise from the specific data set used for training. Thus, the performance of supervised ML tools is only as good as the data used to train them.

Recent works that leverage supervised ML for model parameter estimation typically employ one of two training strategies: (1) parameter combinations obtained from traditional model fitting and the corresponding measured qMRI signals [4] [6] [9] [14] [15] [11] [16] [17], or (2) parameters sampled uniformly from the entire plausible parameter space with simulated qMRI signals [5] [18] [19] [20] [21] [22] [23] [24]. While both of these approaches are limited by the model used to estimate parameters or simulate signals, simulations allow considerably more freedom in choosing training data [25] [26] [27]. However, it is not clear how best to utilise this freedom, as the impact of training data distribution on parameter estimation has yet to be examined.

In this work, we focus on dMRI as an exemplar case and investigate the effect of training data distribution on microstructural parameter estimates. To this end, we quantify bias and variance in estimates throughout the parameter space of a simple dMRI model where the complexity of the estimation task and the dimensionality of the parameter space are low. Specifically, we use a simple two-compartment model based on the spherical mean technique (SMT) [28] [29], which has only two independent parameters. We estimate the microstructural parameters of this model using both traditional non-linear optimisation and supervised ML trained on different distributions of ground truth parameters. We visualise how bias and variance manifest throughout the parameter space, and how regions of high and low estimation performance depend on the distribution and noise level of the training data. Although here we focus on dMRI, we expect similar results and conclusions for other qMRI techniques that use supervised ML methods for fitting multi-compartment models.

## 2. Methods

### 2.1. Data acquisition and pre-processing

After informed written consent, six healthy volunteers were scanned on a 3T Siemens Prisma scanner using a 64-channel head coil. Ethical approval for the study was obtained from the UCL Research Ethics Committee. We acquired diffusion weighted images with b-values of [1000, 2000, 3500, 5000] s/mm^2^ and a total of 128 uniformly distributed gradient directions [30] with 32 gradient directions for each b-value. We acquired 13 b0 images with no diffusion weighting, including one b0 image with reversed phase encoding. Measurement parameters include isotropic 2 mm resolution with acquisition matrix 128 × 128 × 70, partial Fourier imaging 0.75, TE = 94 ms, TR = 9.2 s and GRAPPA parallel imaging with acceleration factor 2. The SNR of the diffusion images was approximately 25 based on the b0 images and averaged across white and grey matter. Additionally, a 3D T1-weighted MPRAGE with 1 mm isotropic resolution was acquired and segmented using FreeSurfer [31] to identify white and grey matter regions in the brain.

To pre-process the diffusion data, we first removed Gibbs ringing artefacts using the method described in [32]. Using the FSL toolbox [33], we estimated the susceptibility-induced off-resonance field with two b0 images with reversed phase encoding polarities [34] and corrected for susceptibility and eddy-current induced geometric distortions and subject motion with methods described in [35]. Finally, we created a binary mask to remove non-brain regions [36].

### 2.2. Biophysical Model

In this work, we use the two-compartment SMT model [28] [29] as a convenient example model that consists of only two independent parameters, which makes visualisation of the parameter space straightforward. In this model, brain tissue is assumed to consist of heterogeneously oriented cylindrical compartments and the surrounding extra-cellular volume. The model can be summarised as

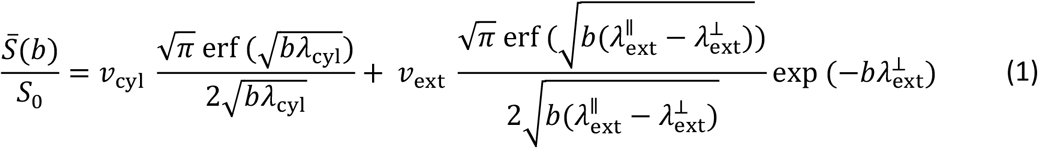

where erf is the error function such that 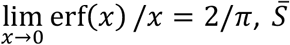, is the powder-averaged diffusion signal at a specific b-value (*b*), *S*_0_ is the signal with no diffusion weighting, *v*_cyl_ and *v*_ext_ are the cylindrical and extra-cellular volume fractions, respectively, *λ*_cyl_ is the diffusivity parallel to cylindrical compartments, and 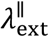 and 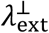 are the parallel and perpendicular extra-cellular diffusivities, respectively. The model assumes that within cylindrical compartments, perpendicular diffusivity is negligible, i.e.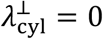, that *v*_cyl_ + *v*_ext_ = 1, and that the extra-cellular diffusivities may be approximated by a tortuosity approximation [37], whereby 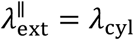 and 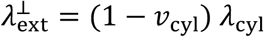. Thus, the model has two independent parameters: *v*_cyl_ and *λ*_cyl_.

### 2.3. Parameter estimation

We estimate the parameters of the biophysical model using two methods: (1) traditional model fitting that utilises non-linear least squares optimisation (software available at https://github.com/ekaden/smt) and (2) supervised ML consisting of artificial neural networks implemented using TensorFlow 2.0 (https://www.tensorflow.org), as well as random forest regressors implemented in Scikit-learn [38]. The following subsections detail the properties of the artificial neural networks, the random forest regressors and the training data.

#### 2.3.1. Artificial neural network architecture

The inputs to the artificial neural networks are the powder-averaged and T2-normalised diffusion signals for the four b-values used: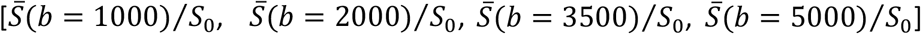. The networks consist of fully connected layers with rectified linear unit (ReLU) activation functions. We include three fully connected layers (input layer, 1 hidden layer, output layer) for the artificial neural networks trained with noise and nine layers (input layer, 7 hidden layers, output layer) for the artificial neural networks trained without noise (i.e. infinite SNR), as more learning capacity is needed to map parameters to noise-free data. Each hidden layer contains 280 nodes. For training, we use a stochastic gradient descent optimiser with learning rate=0.001, momentum 0.9 and the mean squared error loss between the predicted and ground truth model parameter values. Each network was trained over 100,000 epochs.

To train the neural networks, we simulated the powder-averaged and T2-normalised diffusion signal, 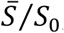, using Equation (1) for each b-value used in this work. Equation (1) provides one signal per b-value, whereas the in-vivo data has 32, one for each gradient direction. Here, we set all 32 measurements in the same b-shell to the same value. We then added noise from a Gaussian distribution with a fixed standard deviation corresponding to a specific SNR. Subsequently, we computed the mean, or powder average, of the noised signals for each b-value. We implemented the noise addition and powder averaging as pre-processing layers in the neural network, as this ensures that a different instance of Gaussian noise is added at each epoch, which in turn ensures that the neural network does not overfit to the noise. In this work, we trained neural networks with three different noise levels corresponding to SNR= [5, 25, ∞].

The neural network outputs are logit(*v*_cyl_) and logit(*λ*_cyl_/*λ*_free_), where logit(x) = log(x) – log(1-x) and *λ*_free_ is the diffusivity of free water, set to 3 μm^2^/ms in this work. The form of the outputs ensures that the parameter estimates lie within a biophysically plausible range, such that 0 ≤ *v*_cyl_ ≤ 1 and 0 ≤ *λ*_cyl_ ≤ *λ*_free_.

#### 2.3.2. Random forest regressor

For the random forest estimator, we used the random forest regressor implemented in Scikit-learn [38] with 200 trees and a maximum tree depth of 20, similarly to previous works [18] [21]. We added noise to the training data and computed the powder average explicitly before training each random forest regressor. The inputs to the random forest regressors are the powder-averaged, T2-normalised signals,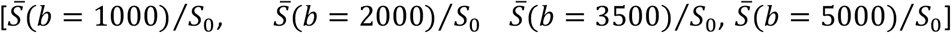, whereas the outputs are logit(*v*_cyl_) and logit(*λ*_cyl_/*λ*_free_), as in the artificial neural network.

#### 2.3.3. Training data distributions

The ML models were trained on synthetic data simulated using Equation (1) and the same set of b-values as in the in-vivo data described in Section 2.1. For each estimator, 2^19^ parameter combinations were drawn from the parameter space bounded by 0 ≤ *v*_cyl_ ≤ 1 and 0 ≤ *λ*_cyl_ ≤ 3 μm^2^/ms, of which 75% were used for training and 25% for validation. We use the following distributions to draw samples for training:

##### (i) Uniform distribution

*v*_cyl_ drawn uniformly between [0, 1], and *λ*_cyl_ drawn uniformly between [0, 3] μm^2^/ms. This distribution corresponds to one of the two approaches used in recent works that estimate tissue microstructure with supervised ML.

##### (ii) Healthy brain distribution

*v*_cyl_ and *λ*_cyl_ sampled using parameter combinations obtained from traditional model fitting in five healthy adult subjects. We fit each of the five healthy adult data sets with traditional model fitting and pooled the resulting parameter combinations. The total number of parameter combinations was approximately 135,000, which is less than the 2^19^ training data samples used in this work. Thus, to ensure that there were sufficient unique parameter combinations for training, we sampled proportionally to the density of parameter combinations obtained from traditional model fitting. First, we computed the two-dimensional histogram of available parameter combinations using 500 bins in both dimensions and used cubic interpolation to approximate the continuous density function *d*(*v*_cyl_, *λ*_cyl_) throughout the *v*_cyl_ - *λ*_cyl_ parameter space. We then performed rejection sampling by selecting a random sample *d’* between the minimum and maximum of the density, as well as a random parameter combination *v*_cyl_’ and *λ*_cyl_’. We computed *d*(*v*_cyl_’, *λ*_cyl_’), and if *d’* < *d*(*v*_cyl_’, *λ*_cyl_’), the parameter combination was accepted, otherwise it was rejected.

This distribution is an approximation of the second approach used in recent works, whereby ML models are trained on parameter combinations estimated via traditional model fitting and the corresponding measured signals. We make one necessary change which is to simulate the diffusion signals using Equation (1) instead of using the measured signals. This allows for increased flexibility in injecting noise into the training data.

##### (iii) Mixed uniform and healthy brain distribution

half the samples drawn from (i) and half drawn from (ii).

To investigate extreme cases where we train on only white or grey matter parameter combinations, we test two further training data distributions:

##### (iv) Healthy WM distribution

*v*_cyl_ and *λ*_cyl_ sampled similarly as in (ii), but for white matter voxels only, determined from the FreeSurfer [31] segmentations.

##### (v) Healthy GM distribution

*v*_cyl_ and *λ*_cyl_ sampled similarly as in (ii), but for grey matter voxels only, determined from the FreeSurfer [31] segmentations.

#### 2.3.4. Summary of trained ML models

Table 1 summarises the ML estimators trained in this work, as well as the names we use to refer to each estimator in the Results and Discussion sections.

**Table 1.** Summary of the ML models trained in this work indicating whether we used the artificial neural network or the random forest regressor, the training data distribution and noise levels used in each trained model.

| Estimator name | ML model | Training data distribution | SNR of training data |
| --- | --- | --- | --- |
| Net-uniform-SNRINF | Artificial neural network | Uniform distribution | ∞ |
| Net-uniform-SNR25 | Artificial neural network | Uniform distribution | 25 |
| Net-uniform-SNR5 | Artificial neural network | Uniform distribution | 5 |
| Net-healthy-brain-SNRINF | Artificial neural network | Healthy brain distribution | ∞ |
| Net-healthy-brain-SNR25 | Artificial neural network | Healthy brain distribution | 25 |
| Net-healthy-brain-SNR5 | Artificial neural network | Healthy brain distribution | 5 |
| Net-healthy-WM-SNR25 | Artificial neural network | Healthy WM distribution | 25 |
| Net-healthy-GM-SNR25 | Artificial neural network | Healthy GM distribution | 25 |
| Net-mixed-SNR25 | Artificial neural network | Mixed uniform and healthy brain distribution | 25 |
| Net-mixed-SNR5 | Artificial neural network | Mixed uniform and healthy brain distribution | 5 |
| RF-uniform-SNRINF | Random forest regressor | Uniform distribution | ∞ |
| RF-uniform-SNR25 | Random forest regressor | Uniform distribution | 25 |
| RF-uniform-SNR5 | Random forest regressor | Uniform distribution | 5 |
| RF-healthy-brain-SNRINF | Random forest regressor | Healthy brain distribution | ∞ |
| RF-healthy-brain-SNR25 | Random forest regressor | Healthy brain distribution | 25 |
| RF-healthy-brain-SNR5 | Random forest regressor | Healthy brain distribution | 5 |
| RF-mixed-SNR25 | Random forest regressor | Mixed uniform and healthy brain distribution | 25 |

### 2.4. Test data

We tested the impact of the training strategy on (i) in-vivo parameter maps, (ii) the bias and variance of predicted model parameter across the entire parameter space, (iii) the performance of parameter estimation for normal and abnormal parameter combinations, and (iv) the detectability of regions of abnormal tissue in parameter estimates. We outline the data sets used for these four test cases in the following subsections.

#### 2.4.1. In-vivo test data

To compare parameter estimates obtained in healthy human brain scans, we used the diffusion measurements of the 6^th^ healthy adult volunteer that was not included in the training data parameter pool used in distributions (ii)-(v) described in Section 2.3.3. The SNR of this data set was approximately 25, and the images were pre-processed as described in Section 2.1.

#### 2.4.2. Simulated data for different parameter combinations

To probe the overall accuracy and precision of the model fitting, we synthesized test data using Equation 1 with the same set of b-values as in the in-vivo data described in Section 2.1. We chose 441 points on a 21×21 grid covering the parameter space, such that *v*_cyl_ ranged from 0 to 1 at increments of 0.05, and *λ*_cyl_ ranged from 0 to 3 μm^2^/ms at increments of 0.15 μm^2^/ms. For each of the parameter combinations on this grid, we synthesised 10,000 samples of the diffusion signals and added Gaussian noise. We created three such data sets with SNR = [5, 25, ∞]. For each of the test data sets we used the neural networks trained with the corresponding noise level to estimate parameters.

#### 2.4.3. Simulated normal and abnormal parameter combinations

In addition to the gridded parameter combinations in the previous section, we synthesised the diffusion signals for five further parameter combinations to probe specific normal and abnormal tissues (Table 2). These included the mean parameter combination found in white matter based on traditional model fitting for 5 healthy adult subjects (WM), the mean parameter combination found in grey matter based on traditional model fitting for 5 healthy adult subjects (GM), two extreme abnormalities (Abnormality 1 and Abnormality 2), and an abnormality where both *v*_cyl_ and *λ*_cyl_ deviate from WM only slightly (Abnormality 3). For each of these parameter combinations, we synthesised 10,000 samples of the diffusion signals and added Gaussian noise corresponding to SNR = [5, 25]. As before, for each of the test data sets we used the neural networks trained with the corresponding noise level to estimate parameters.

**Table 2.** Specific parameter combinations chosen to illustrate performance in typical and abnormal parameter combinations.

| Parameter combination name | $v_{cyl}$ | $\lambda_{cyl} (\mu\text{m}^2/\text{ms})$ |
| --- | --- | --- |
| Typical white matter (WM) | 0.67 | 2.20 |
| Typical grey matter (GM) | 0.16 | 1.14 |
| Abnormality 1 | 0.67 | 0.50 |
| Abnormality 2 | 0.05 | 2.20 |
| Abnormality 3 | 0.60 | 1.80 |

#### 2.4.4. Simulated brain data with abnormality

Taking the in-vivo parameter maps obtained from traditional model fitting, we chose a region of interest (ROI) in white matter and changed the parameter combinations in this region to those of Abnormality 3. Using the parameter combinations from the in-vivo parameter maps with the altered ROI, we simulated diffusion signals with Equation (1) to create a full simulated brain-like data set. We added noise to the simulated signals corresponding to SNR = [5, 25]. These two noised data sets were used to investigate whether small abnormalities can be visually detected with the different estimation approaches.

## 3. Results

In this section we present the accuracy and precision of parameter estimates using traditional model fitting and ML. For the ML approach, we focus on artificial neural networks as an example but obtain similar results using the random forest regressors.

### 3.1. In-vivo parameter maps

We map in-vivo parameter estimates for a single healthy adult subject using traditional model fitting (Figure 1A) and using Net-uniform-SNR25, Net-healthy-brain-SNR25, Net-healthy-WM-SNR25 and Net-healthy-GM-SNR25 (Figure 1B). Figure 1 demonstrates that when we train only on parameter combinations typical in white matter, estimates in grey matter are substantially different from those obtained from traditional model fitting, whereas when we train only on parameter combinations typical in grey matter, estimates in white matter are substantially different from those obtained from traditional model fitting. Parameter maps obtained using Net-uniform-SNR25 and Net-healthy-brain-SNR25 are comparable to those from traditional model fitting.

**Figure 1.**
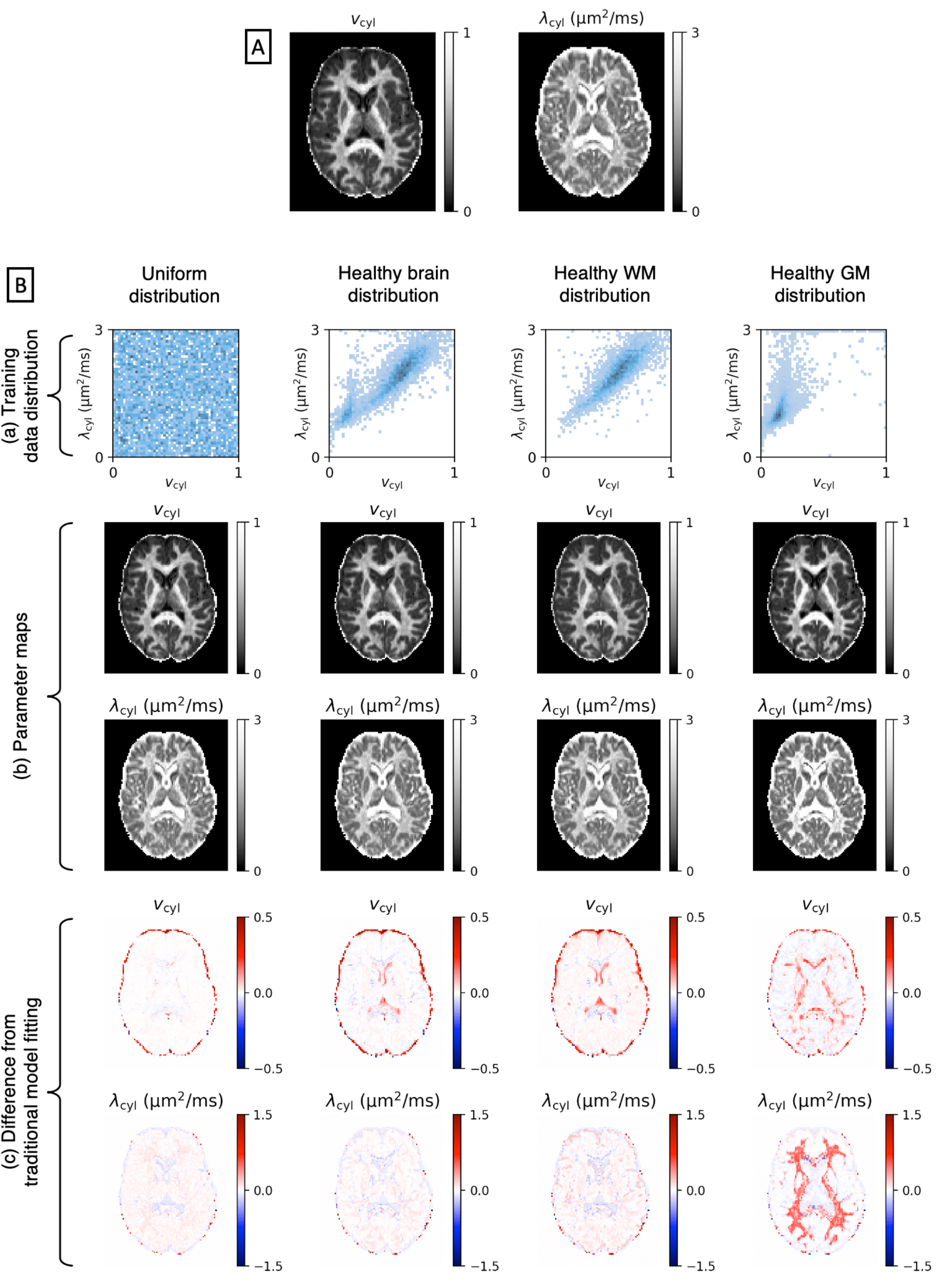
Panel (A): v_cyl_ and λ_cyl_ parameter maps obtained from traditional model fitting. Panel (B): (a) Different training data distribution strategies, (b) the corresponding v_cyl_ and λ_cyl_ parameter maps and (c) the difference between parameter maps in row (b) and parameter maps from traditional model fitting in Panel (A).

### 3.2. Accuracy and precision using synthetic test data

In this section, we use synthetic test data to compare the accuracy and precision of parameter estimates obtained using traditional model fitting and artificial neural networks trained on different data distributions at different noise levels.

Figure 2 maps bias in parameter estimation for different combinations of *v*_cyl_ and *λ*_cyl_. The arrows point from the ground truth parameters to the mean of estimated parameters. Different rows show the different noise levels that were injected to both the training data and the test data. As SNR is reduced, bias in the parameter estimates increases for each estimation method, with traditional model fitting providing the lowest overall bias. Estimates obtained from the artificial neural network trained on the healthy brain distribution has the highest overall bias, and bias is consistently high in the low *v*_cyl_ and high *λ*_cyl_ region where the training data has low density. Interestingly, certain regions of the parameter space act as ‘sinks’, towards which estimates of nearby parameters are biased. The location of these sinks depends on both the training data distribution and the noise level. For example, in the networks trained on in-vivo parameter combinations a sink forms near the highest data density region. The pull of the sink becomes stronger as the SNR is reduced. For each fitting approach, biases are high when *λ*_cyl_ = 0, as the biophysical model is degenerate when there is no diffusion. We obtained similar results using random forest regressors (see Supplementary Figure S1A).

**Figure 2.**
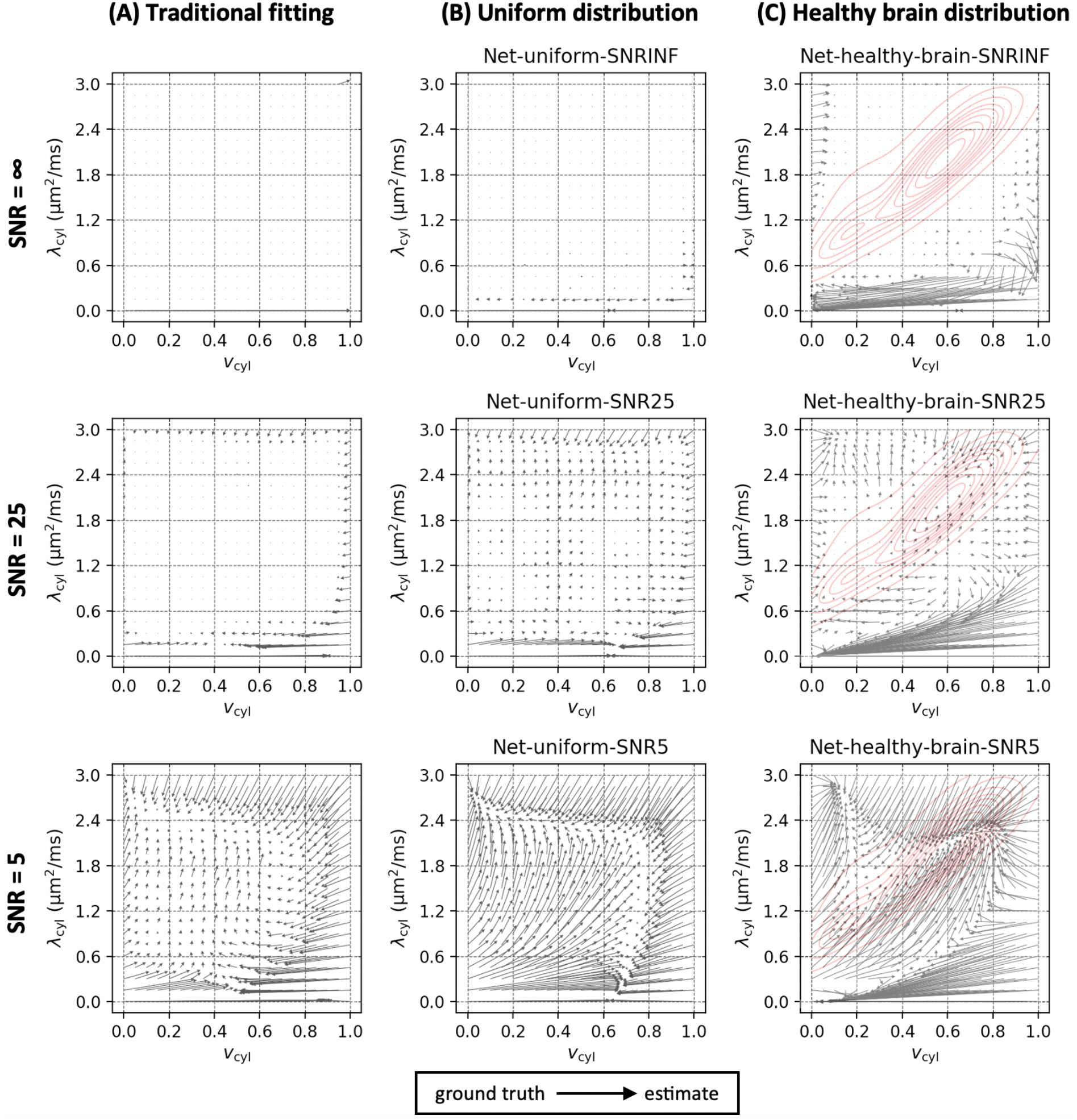
Bias mapped using quiver plots for (A) traditional model fitting, (B) neural networks trained using the uniform distribution and (C) neural networks trained using the healthy brain distribution. The arrows point from the ground truth values to the mean of the estimated values. In column (C), the red contours show the training data density. Each row shows the biases at different values of SNR, according to which Gaussian noise was added both the training data and test data.

Figure 3 shows the standard deviation in *v*_cyl_ and *λ*_cyl_ estimates obtained from traditional model fitting and from the artificial neural networks. Parameters are estimated precisely using all three methods when the training and test data are noise-free. As SNR is reduced, the precision of the parameter estimates obtained using traditional model fitting degrades more than using the artificial neural networks. We obtained similar results using random forest regressors (see Supplementary Figure S1B).

**Figure 3.**
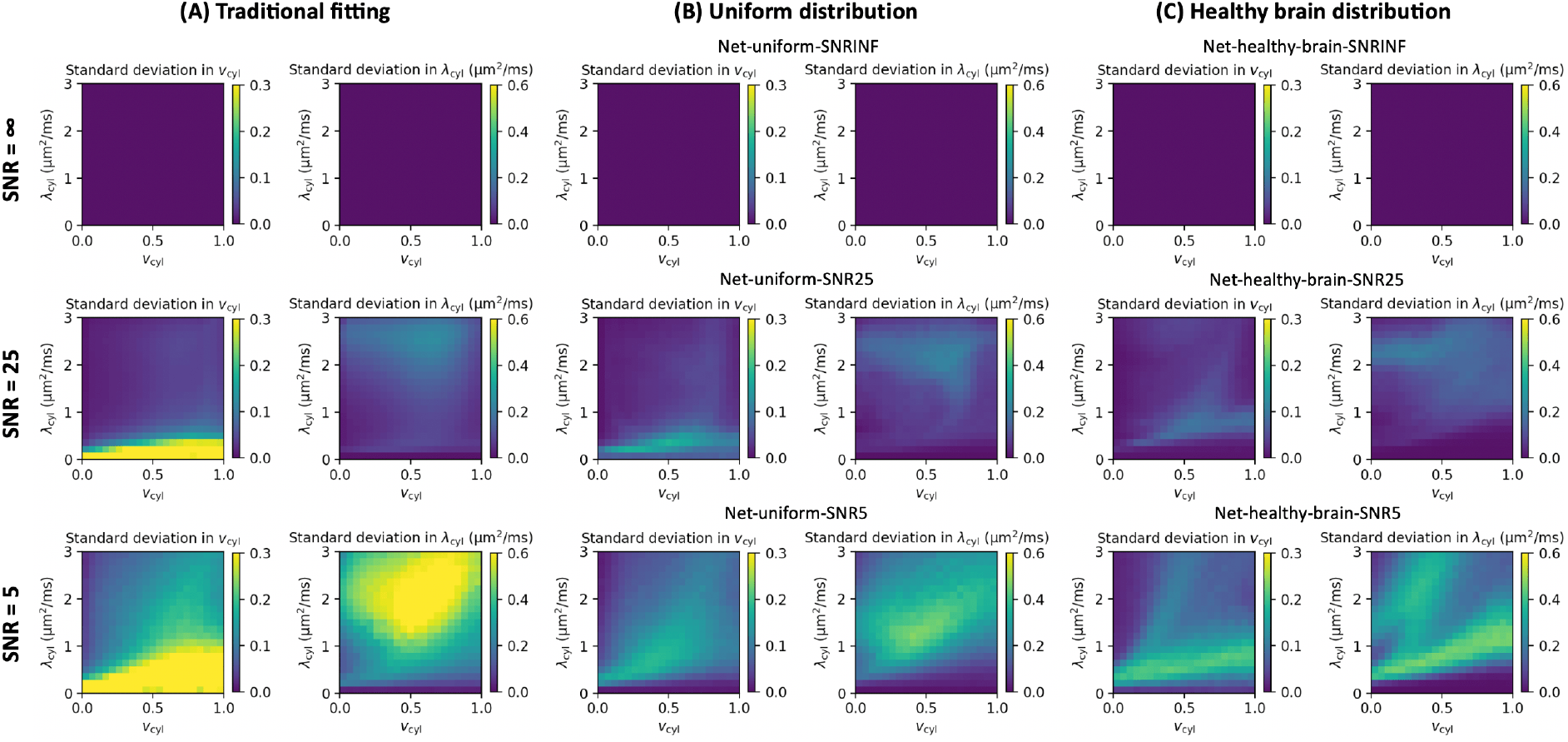
Precision of v_cyl_ and λ_cyl_ estimates using (A) traditional model fitting, (B) neural networks trained using the uniform distribution and (C) neural networks trained using the healthy brain distribution. The three rows correspond to the different noise levels in both the training and test data sets.

In Figure 4, we probe estimation performance for the specific parameter combinations representing white matter (WM), grey matter (GM) and three tissue abnormalities outlined in Section 2.4.2. We compare traditional model fitting and the neural networks trained on uniform, healthy brain and mixed uniform-healthy brain distributions. When SNR = 25, WM and GM are estimated accurately using all estimation methods. Precision is comparable across the estimation methods for GM, but precision in WM estimates obtained using Net-uniform-SNR25 is slightly lower than using Net-mixed-SNR25 and Net-healthy-brain-SNR25. When SNR = 5, biases appear in WM and GM estimates obtained using the neural networks. In WM, precision is low using traditional model fitting compared to the neural networks, whereas in GM, precision is lowest using Net-uniform-SNR5.

**Figure 4.**
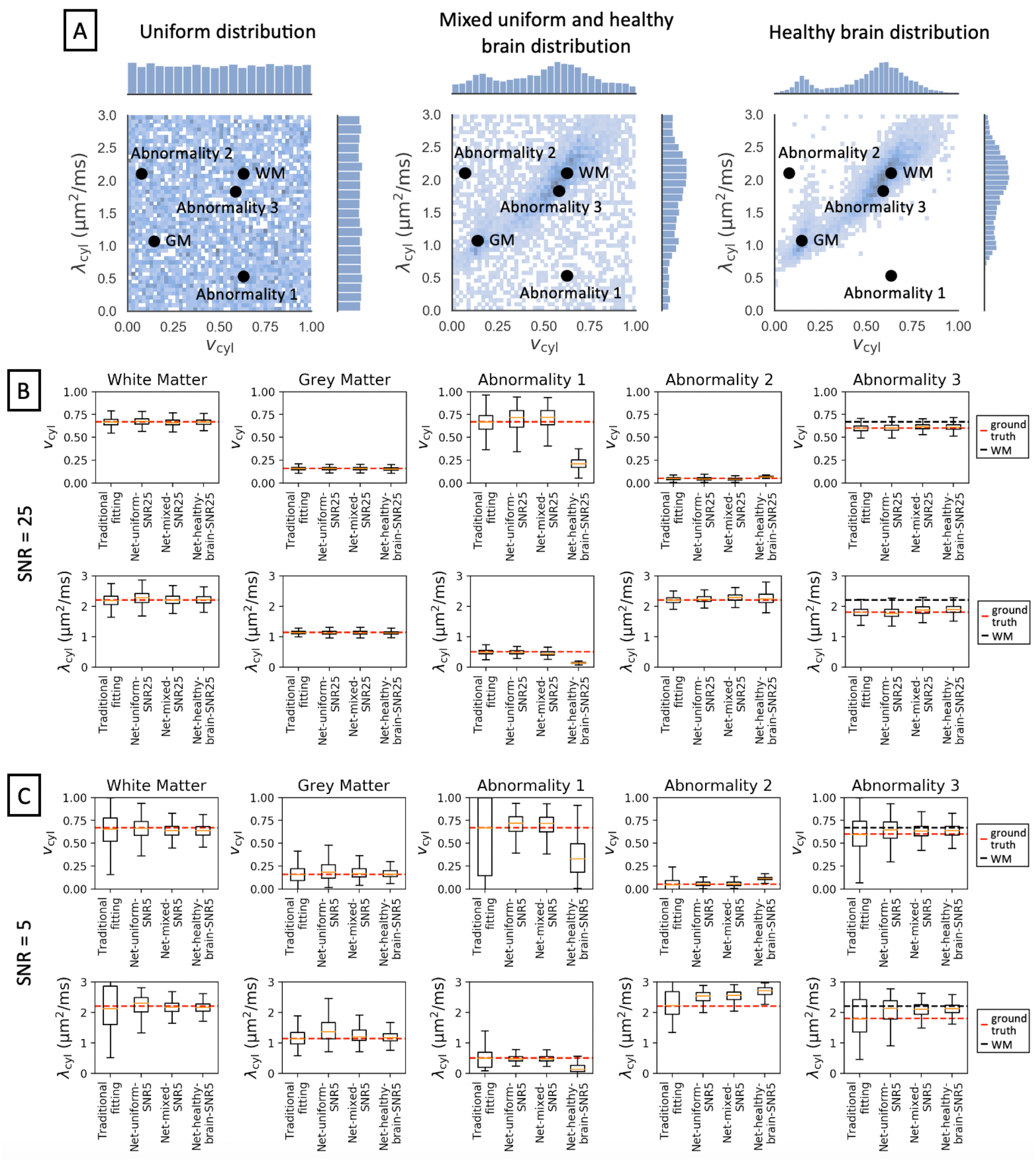
Panel (A): Different training data distributions: uniform data distribution, healthy brain distribution, and a mixed distribution where 50% of the samples are from the uniform distribution, and 50% of the samples are from the healthy brain distribution. We mark five parameter combinations: white matter (WM), grey matter (GM), two different extreme parameter combinations (Abnormality 1 and 2) and one parameter combination that differs only slightly from typical WM (Abnormality 3). We show box plots of the estimates for these five parameter combinations using synthetic data with SNR = 25 in panel (B) and using synthetic data with SNR = 5 in panel (C). The dashed red line marks the ground truth, and for Abnormality 3, the black line marks normal WM.

Abnormality 1 is estimated with low accuracy using the neural networks trained on both the healthy brain distribution and mixed uniform and healthy brain distribution. Biases are substantial when SNR = 25 and are exacerbated for SNR = 5. For SNR = 25, estimates of Abnormality 2 are biased using Net-healthy-brain-SNR25, and as SNR is decreased to 5, estimates of Abnormality 2 are biased using all three neural networks. For Abnormality 3, estimates tend to be accurate using all methods when SNR = 25. However, as SNR is decreased to 5, the estimates using the neural networks are biased toward WM values. We demonstrate this effect further in Figure 5 using synthetic brain-like test data described in Section 2.4.3. When SNR = 25, Abnormality 3 can be visually distinguished from surrounding healthy tissue for all the estimation methods. When SNR = 5, estimates from traditional model fitting are noisy throughout the brain, whereas estimates from the neural networks appear smooth, particularly for Net-mixed-SNR5 and Net-healthy-brain-SNR5, but Abnormality 3 cannot easily be distinguished from the surrounding tissue.

**Figure 5.**
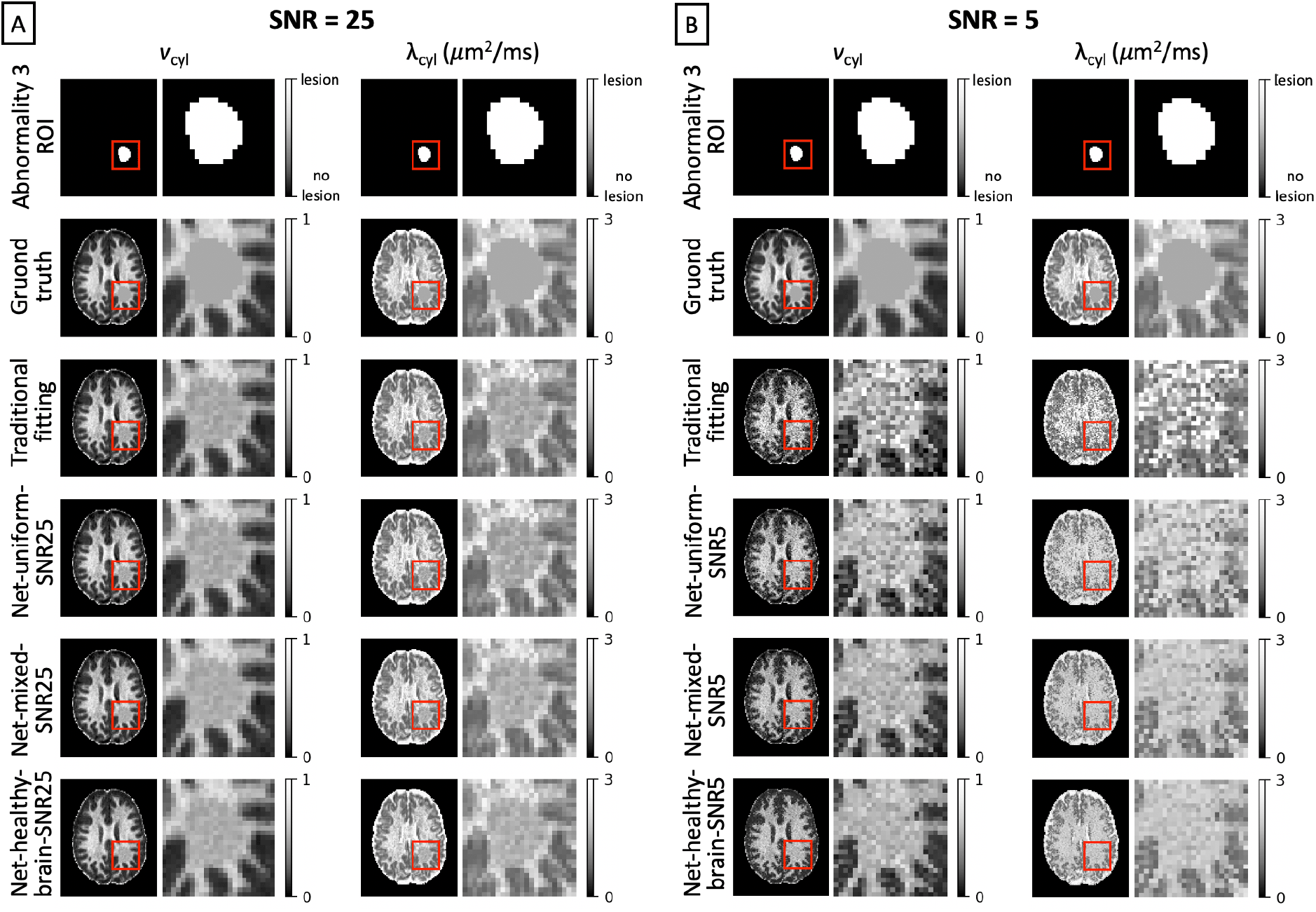
Parameter estimates for (A) SNR = 25 and (B) SNR = 5. The data sets used here were simulated using parameter values obtained from traditional fitting with Abnormality 3 applied to an ROI shown in the top row. Abnormality 3 is highlighted in the red box and shown in adjacent zoomed plots.

### 3.3. Summary of results

In Table 3 we summarise the overall RMSE, bias and standard deviation using the different parameter estimation methods for SNR = 25. Net-uniform-SNR25 yields the lowest average RMSE for the estimation methods tested in this work. On average, traditional fitting yields the lowest average bias in both *v*_cyl_ and λ_cyl_ with Net-uniform-SNR25 in second place. For *v*_cyl_, bias is approximately 8% higher using Net-uniform-SNR25 compared to traditional model fitting, whereas for λ_cyl_, bias is almost 200% higher using Net-uniform-SNR25 compared to traditional model fitting. Net-healthy-brain-SNR25 and RF-healthy-brain-SNR25 yield the lowest standard deviations in *v*_cyl_ and λ_cyl_, respectively, but at the cost of high average biases in both parameters compared to the other estimation methods.

**Table 3.**
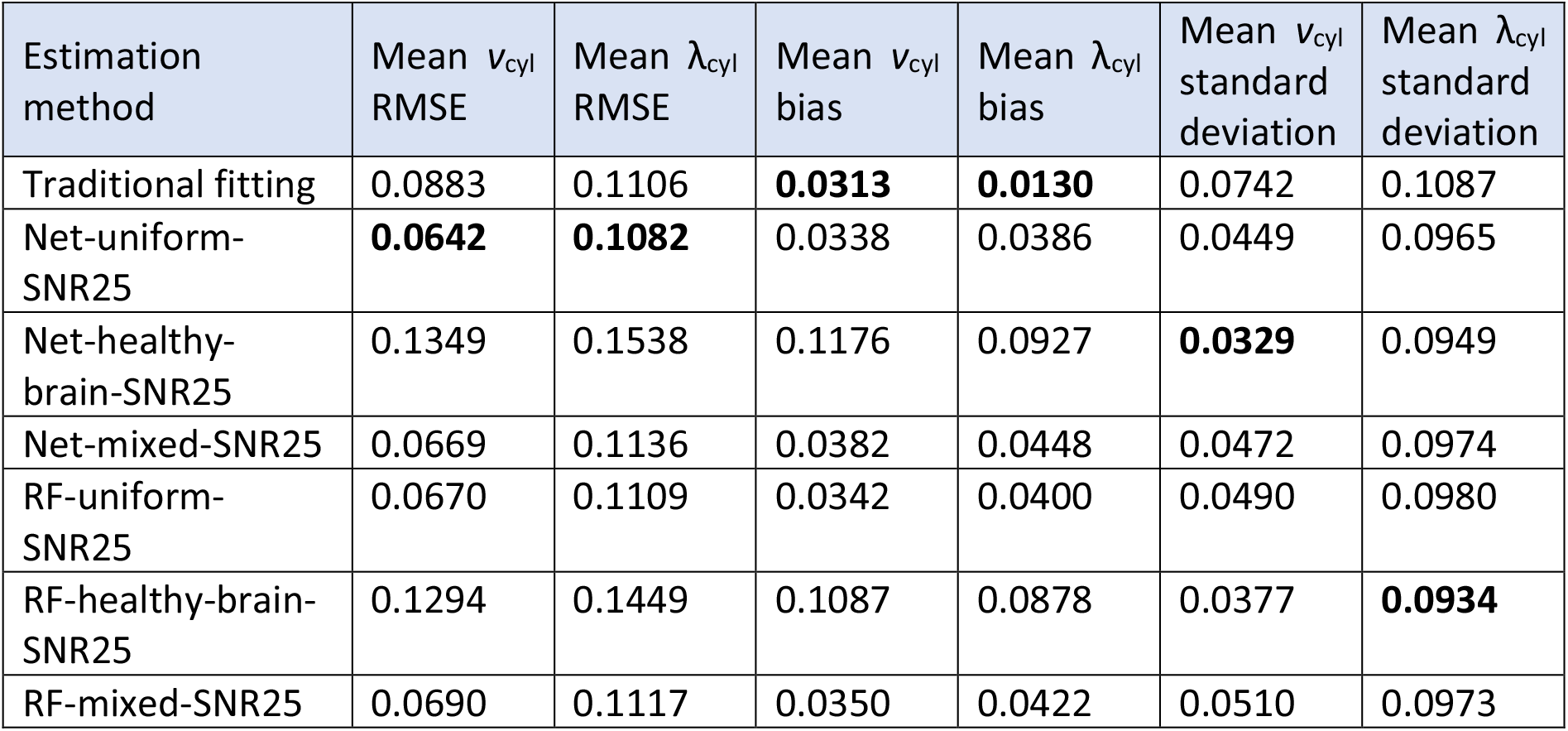
The mean RMSE, bias and standard deviation over the entire parameter space v_cyl_ and λ_cyl_ for estimation methods using SNR = 25. Bold values highlight the lowest value in each column.

| Estimation method | Mean $v_{cyl}$ RMSE | Mean $\lambda_{cyl}$ RMSE | Mean $v_{cyl}$ bias | Mean $\lambda_{cyl}$ bias | Mean $v_{cyl}$ standard deviation | Mean $\lambda_{cyl}$ standard deviation |
| --- | --- | --- | --- | --- | --- | --- |
| Traditional fitting | 0.0883 | 0.1106 | **0.0313** | **0.0130** | 0.0742 | 0.1087 |
| Net-uniform-SNR25 | **0.0642** | **0.1082** | 0.0338 | 0.0386 | 0.0449 | 0.0965 |
| Net-healthy-brain-SNR25 | 0.1349 | 0.1538 | 0.1176 | 0.0927 | **0.0329** | 0.0949 |
| Net-mixed-SNR25 | 0.0669 | 0.1136 | 0.0382 | 0.0448 | 0.0472 | 0.0974 |
| RF-uniform-SNR25 | 0.0670 | 0.1109 | 0.0342 | 0.0400 | 0.0490 | 0.0980 |
| RF-healthy-brain-SNR25 | 0.1294 | 0.1449 | 0.1087 | 0.0878 | 0.0377 | **0.0934** |
| RF-mixed-SNR25 | 0.0690 | 0.1117 | 0.0350 | 0.0422 | 0.0510 | 0.0973 |

## 4. Discussion

This work highlights two key properties of supervised ML-based fitting techniques, which differ from traditional model fitting. Firstly, we show that parameter estimates are significantly affected by the distribution of training data. Secondly, we demonstrate that smooth parameter maps obtained via ML may be deceptive, as high precision may hide strong biases. This is in contrast with traditional fitting, where low reliability in estimates is typically reflected by noisy parameter maps. The results presented in this work focus on artificial neural networks as the example for supervised ML, but we observe similar trends with other ML models such as random forest regressors, for which we summarise accuracy and precision in Supplementary Figure S1.

In Section 3.2. we focus on three different training data distributions: healthy parameter combinations obtained using traditional model fitting, uniformly distributed parameter combinations, and healthy parameter combinations augmented with uniformly distributed parameter combinations. Recently, authors in [39] compared the fitting performance of the first two training strategies, and authors in [40] assessed the trade-off between accuracy and generalisability when combining them to analyse diffusion-relaxation data. Our results show that training on healthy parameter combinations facilitates precise estimates in healthy tissue but may yield strong biases in atypical parameter combinations not represented in training.

This bias is mitigated when healthy data is combined with atypical parameter combinations in training, in line with recent findings in [40]. However, here we show that even when healthy training data is combined with atypical parameter combinations, and in fact even when the full parameter space is uniformly represented in the training data, supervised ML may still introduce substantial biases that can hamper the clinical utility of qMRI techniques. Thus, our findings suggest that it is crucial to develop training strategies that minimise biases throughout the parameter space.

Parameter estimates obtained from traditional model fitting are overall more accurate than the estimates obtained from the ML models at each noise level tested in this work. However, at low SNR traditional fitting suffers from high variance, which limits the interpretability of estimated parameter maps. Maps obtained using the neural networks are less noisy, which may mistakenly convince the user that the estimates are reliable even at low SNR. Indeed, in Figure 6 we show that a small white matter abnormality may be missed if the low SNR neural network estimates are trusted. While this issue is particularly pronounced for the networks trained on healthy parameter combinations, the maps obtained using the uniform distribution may mislead users as well. Our findings highlight the importance of accounting for bias and variance of model parameter estimates when using supervised ML methods for model fitting tasks. The analysis and visualization approaches proposed here (Figures 2-5) provide tools to quantify the expected impact of a chosen estimation strategy and to aid the interpretation of resulting parameter estimates. For example, parameter estimates near ‘sinks’ in the bias quiver plots should be interpreted with caution, as these parameter combinations may mask substantial biases. The location and evolution of these sinks can inform future experimental design and training strategies optimised to mitigate their impact. Furthermore, our findings point out the importance of computing uncertainty, cf. [41], in ML-based estimation, particularly when ML is used to compensate for lower quality data.

This work used a simple two-compartment model to demonstrate the impact of training data on a low-dimension system in dMRI. We expect to see similar effects in other models and qMRI techniques, likely exacerbated by complexity, but verification of this will be the subject of future work. Our analysis was also limited to a single set of b-values. Different numbers and combinations of b-values would likely affect both the overall accuracy and the position of ‘sinks’ in the parameter space towards which nearby parameter combinations are biased. Finally, we note that the distribution of noise used to inject both the training and synthetic test data sets is Gaussian, and not Rician, as one might expect from magnitude MRI data. This likely has an impact on the in-vivo estimates obtained in Section 3.1., which were not corrected for Rician noise bias. Future work could investigate how the form, and not just the width of the noise distribution may affect parameter estimates in in-vivo measurements.

ML is a promising tool for enhancing medical imaging technology, where resources are often limited, and the potential impact may be life changing. qMRI may benefit in particular, as advanced MRI acquisitions and subsequent model fitting may be time-consuming. However, work still needs to be done to mitigate biases and assess estimation reliability in order to use ML effectively. Here, we use a two-compartment model and dMRI data to highlight with a simple example that performance depends strongly on the choice of training data. Future work might explore optimising training data sampling given a set of experimental parameters and tissue model.

## Acknowledgements

NGG thanks the London Interdisciplinary Bioscience PhD Consortium and is funded by BBSRC grant BB/M009513/1. MP is supported by UKRI Future Leaders Fellowship (MR/T020296/1). EPSRC grants EP/M020533/1 and EP/N018702/1 and the NIHR UCLH Biomedical Research Centre and the NIHR GOSH Biomedical Research Centre support our work on this topic. We acknowledge Enrico Kaden’s background contributions in conceptualisation, design and set-up of the diffusion experiment, and initial discussions on studying the effects of training data distribution. We also thank Filip Szczepankiewicz and Markus Nilsson for sharing their diffusion EPI sequence.

## Supplementary Material

**Figure S1.**
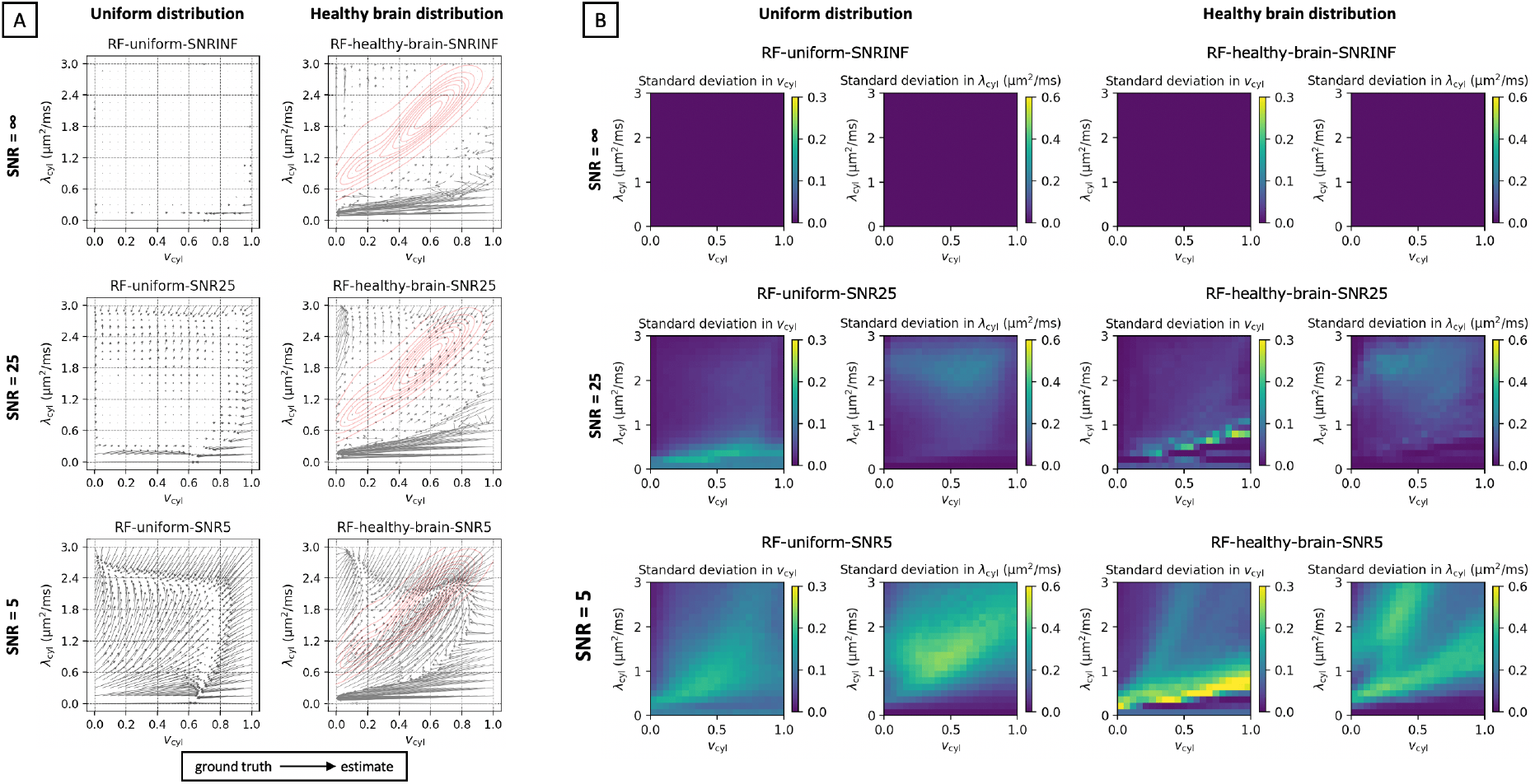
Estimation performance using a random forest regressor. Panel (A) shows biases and panel (B) shows the standard deviation for different noise levels and training data distributions using synthetic data.

## Notes

### Competing Interest Statement

The authors have declared no competing interest.

